# A non-dimensional, two-parameter mechanical model reveals alterations in nuclear mechanics upon Hepatitis C Virus infection

**DOI:** 10.1101/395129

**Authors:** Sreenath Balakrishnan, M S Suma, Geetika Sharma, Shilpa R Raju, Uma Reddy, Saumitra Das, G. K. Ananthasuresh

## Abstract

Morphology of the nucleus is an important regulator of gene-expression. Nuclear morphology is in turn a function of the forces acting on it and the mechanical properties of the nuclear envelope. Here, we present a two-parameter, non-dimensional mechanical model of the nucleus that reveals a relationship among nuclear shape parameters such as projected area, surface area and volume. Our model fits the morphology of individual nuclei and predicts the ratio between forces and modulus in each nucleus. We analyzed the changes in nuclear morphology of liver cells due to Hepatitis C Virus (HCV) infection using this model. The model predicted a decrease in the elastic modulus of the nuclear envelope and an increase in the pre-tension in cortical actin as the causes for the change in nuclear morphology. These predictions were validated biomechanically by showing that liver cells expressing HCV proteins possessed enhanced cellular stiffness and reduced nuclear stiffness. Concomitantly, cells expressing HCV proteins showed down-regulation of lamin-A,C and up-regulation of actin, corroborating the predictions of the model. Our modelling assumptions are broadly applicable to adherent, monolayer cell cultures making the model amenable to investigate changes in nuclear mechanics due to other stimuli by merely measuring nuclear morphology. Towards this, we present two techniques, graphical and numerical, to use our model for predicting physical changes in the nucleus.

## Introduction

It is known that cell-function and cell-fate are regulated by mechanical properties of the nucleus (1–3) and its morphological changes brought about by forces acting on it (4). Cancer (5, 6), laminopathies (7–10), and other diseases are known to effect such modifications in nuclei. We present another instance wherein Hepatitis C Virus (HCV) alters nuclear mechanics. We observed that the nuclear size of virus-infected liver cells was more than that of control cells. The underlying mechanism for this was revealed by a two-parameter mechanical model of the nucleus subject to compression by cortical actin and net expansion by osmotic pressure, chromatin, and microtubules. Figure 1 illustrates these two competing forces acting on a nucleus in monolayer cell culture as in (11). As shown in the figure, compressive force by cortical actin can be approximated as planar pressure pushing the nucleus down. This is akin to pressing an inflated sphere with a flat plate. Incidentally, the deformation analysis of such an elastic sphere was reported in (12) using an axi-symmetric model that led to a pair of ordinary differential equations. The solutions to these equations depend only on two non-dimensional parameters: (i) *η*_1_ *= PR*/2*E*_1_*H*, the ratio between the expanding pressure, *P*, to the elastic modulus, *E*_*1*_, of the nuclear envelope of radius, *R*, and thickness, *H*, in the unloaded state (Fig. 1*B*); and (ii) 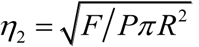, the ratio between the compressive force, *F*, to *P*. In order to estimate *η*_1_ and *η*_2_ corresponding to individual, experimentally measured, nuclear morphological features, we first calculated three parameters to characterize nuclei individually: (i) Projected area (*A*_*p*_), (ii) Surface area (*A*_*s*_) and (iii) Volume (*V*). Next, we normalized these parameters using *R* to define *a_p_ = A_p_/R*^2^, *a_s_ = A_s_/R*^2^ and *v = V/R*^3^. Our model enables expressing *η*_*1*_ and *η*_2_ in terms of *a*_*p*_, *a*_*s*_, and *v*. The three normalized parameters were then used to fit the model and thereby obtain *η*_1_ and *η*_2_. Difference in *η*_1_ and *η*_2_ between control and HCV-infected liver cells suggested changes in *E*_*1*_ and *F* due to HCV. These predictions from the model were experimentally verified using Atomic Force Microscopy (AFM), western blot and immunofluorescence.

**Fig. 1.**
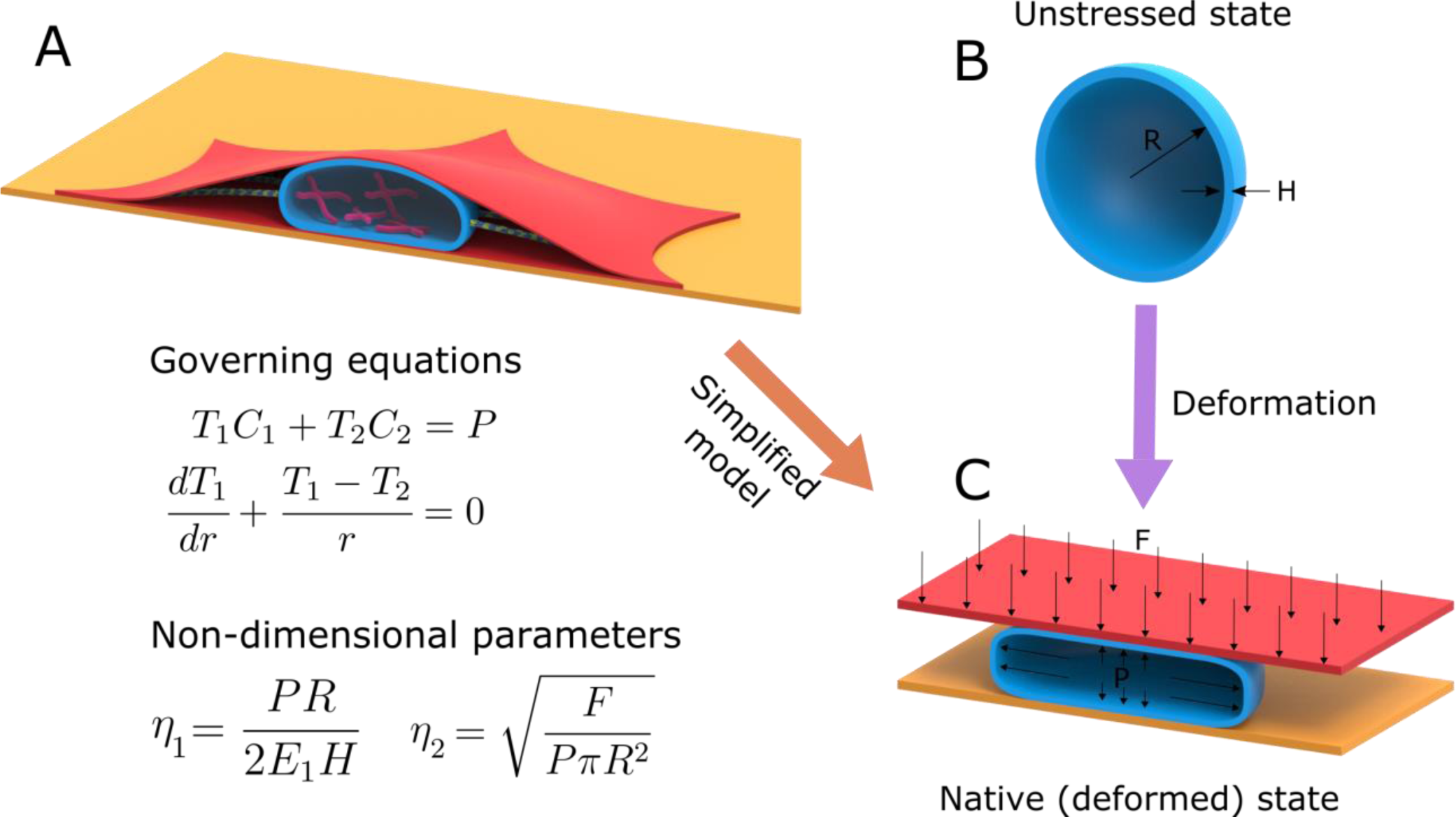
A mechanical model of nuclear morphology – (A) Nuclear envelope (blue) is shaped by forces from the nucleoplasm and cytoplasm. These forces are mainly due to cortical actin (red), microtubules (green), chromatin (pink) and an osmotic pressure difference between the nucleoplasm and cytoplasm. (B) In the absence of forces, the nucleus is assumed to be a spherical membrane of radius *R* and thickness *H*. (C) The net contribution from osmotic pressure, microtubules and chromatin is assumed to be an inflating pressure *P*. The force due to cortical actin, *F*, is assumed to be originating from a flat plate that is pushing down on the nucleus. The equations of equilibrium of the membrane in the normal and tangential directions were obtained from (12) where *T*_*1*_ and *T*_*2*_ are the forces per unit length in the principal directions; and *C*_*1*_ and *C*_*2*_ are the principal curvatures. The solutions to the equations depend only on two non-dimensional parameters *η*_*1*_ and *η*_*2*_.

The changes in *E*_1_ and *F* were estimated using AFM by measuring nuclear and cell stiffness. Nuclear stiffness can be used as an indicator of *E*_*1*_ and cell stiffness as an indicator of *F*. This is because nuclear stiffness increases with expression of lamin-A,C (13), which in turn is known to be a major elastic component of the nuclear envelope (14). Additionally, pre-tension in cortical actin is known to correlate with cell stiffness (15). Differential expression of lamin-A,C and actin measured using western blot and immunofluorescence independently reinforced the predictions from our model as well as AFM measurements.

We used HCV replicon cells (16) that constitutively harbor HCV replicon RNA as a model system with Huh7 cell line as the control. An important biologically relevant finding of this work is that HCV proteins down-regulate lamin-A,C and up-regulate actin. This finding is supported by morphological and biomechanical measurements with the help of the aforementioned two-parameter model. Thus, we show consistency in the results of three different experiments, namely, biochemical assays, AFM stiffness measurements, and morphological parameters.

i. Biochemical assays showed downregulation of lamin-A,C and upregulation of *β* - actin in HCV replicon cells as compared to Huh7 cells. Based on the reported literature, we associate these two changes to reduced elastic modulus of the nuclear membrane and enhanced cortical actin force acting on the nucleus. The two changes correspond to smaller *E*_1_ and larger *F* in replicon cells than those in the control. These changes could be rescued upon over-expressing lamin-A in replicon cells.
ii. Force-displacement characteristics from AFM measurements on replicon and control cells, with and without disruption of actin, were found to be consistent with the observed trends in *E*_1_ and *F*. That is, HCV replicon cells showed higher stiffness than control cells when actin was intact. This supports the increased pretension in cortical actin in HCV replicon cells. On the other hand, upon disruption of actin using Cytochalasin-D, indentation using AFM is indicative of elastic modulus of the nuclear membrane. In this case, replicon cells showed lower stiffness than control cells.
iii. Nuclei of replicon cells have larger volume, surface area, and projected area than control cells. The increase in size of the nucleus of replicon cells was rescued with over-expression of lamin-A.

These experimental results are tied together with the help of a simplified elastic model of the nucleus of an adherent cell. It is worth mentioning that three morphological parameters (*a*_*p*_, *a*_*s*_ and *v*) are adequate to establish relative changes in the elastic modulus of nuclei and force acting on them, which in turn are related to changes in lamin-A,C and actin. Its significance lies in the fact that the volume and area of individual nuclei are easily quantified experimentally and computationally. This simple and direct biomechanical assay of extracting physical properties of nuclei of a large population is an overarching contribution of this work.

## Materials and Methods

### Non-dimensional, mechanical model for the nucleus

Nuclear morphology is a consequence of mechanical equilibrium between the forces acting on the nuclear envelope and the stresses generated inside it. Stresses in the nuclear envelope are in turn a function of the forces acting on it and its mechanical properties. Therefore, information about these forces and mechanical properties is contained in the morphology of the nucleus. For a given shape of the nucleus, we estimated the forces acting on the nuclear envelope and its mechanical properties.

We assumed that in its native state under mechanical equilibrium, there are predominantly two forces acting on the nuclear envelope: (i) a uniform pressure that inflates the nucleus and (ii) a downward force from cortical actin that compress the nucleus (Fig. 1*C*). These forces are qualitatively similar to those in (11). In that study, the authors assumed that the inflating pressure is the net result of the osmotic pressure due to difference in solute concentration between the nucleoplasm, and cytoplasm; and a compressive force from the microtubules. However, some previous studies have argued that the osmotic pressure is due to a difference in concentration of macromolecules, not solutes, across the nuclear envelope (17, 18). In another study, additional forces on the nuclear envelope, due to chromatin, were assumed (19). As the origin of pressure is unimportant in our study, we considered all contributions from the osmotic pressure, microtubules and chromatin into a single inflating pressure.

The force due to cortical actin is assumed to be arising from a flat plate pushing down on the nucleus (Fig. 1*C*) as in (11). Since we considered the steady-state morphology of the nuclear envelope, we neglected the effects due to viscosity. Viscosity would affect the rate of convergence of the changing morphology but not the final morphology (11). The nuclear envelope was hence assumed to be a hyperelastic membrane that is spherical in the unloaded state (11, 19, 20). Its bending was neglected because the nuclear envelope is relatively thin in the order of 100 nm as compared to the size of the nucleus, which is around 10 μm in diameter. We assumed the nuclear envelope to be made of an incompressible Mooney-Rivlin material. Unlike in some previous studies (11, 19), we did not choose a neo-Hookean model since it exhibits instabilities at large strains (21). The instability refers to the drop in pressure at large strains (> 38%) when a spherical, neo-Hookean membrane is inflated (21).

Since the initial geometry, forces and boundary conditions are axisymmetric, we used an analytical formulation developed for mechanical equilibrium of axisymmetric membranes (12). When the governing ordinary differential equations for mechanical equilibrium were expressed in terms of the principal strains (see supplementary information), we obtained the two non-dimensional parameters that govern the deformation, namely, (i) *η*_1_ *= PR/*2*E*_1_*H* and (ii) 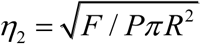. While *η*_1_ appears in the equation for force equilibrium, *η*_2_ appears in the boundary condition (see supplementary information). In order to solve the governing differential equations, we require two other parameters: (i) *λ*_0_, the stretch at the apex point of the nucleus in the deformed state (point M’ in Fig. 3*A*) and (ii) *τ*, half of the angle subtended by the region of contact between cortical actin and nuclear envelope in the undeformed state (Fig. 3*A*). However, as there are only two independent parameters, by specifying either of them, the other two can be determined (see supplementary information). In our simulations we have specified *λ*_0_ and *τ*, and estimated *η*_1_ and *η*_2_ (Fig. 3*F*).

**Fig. 2.**
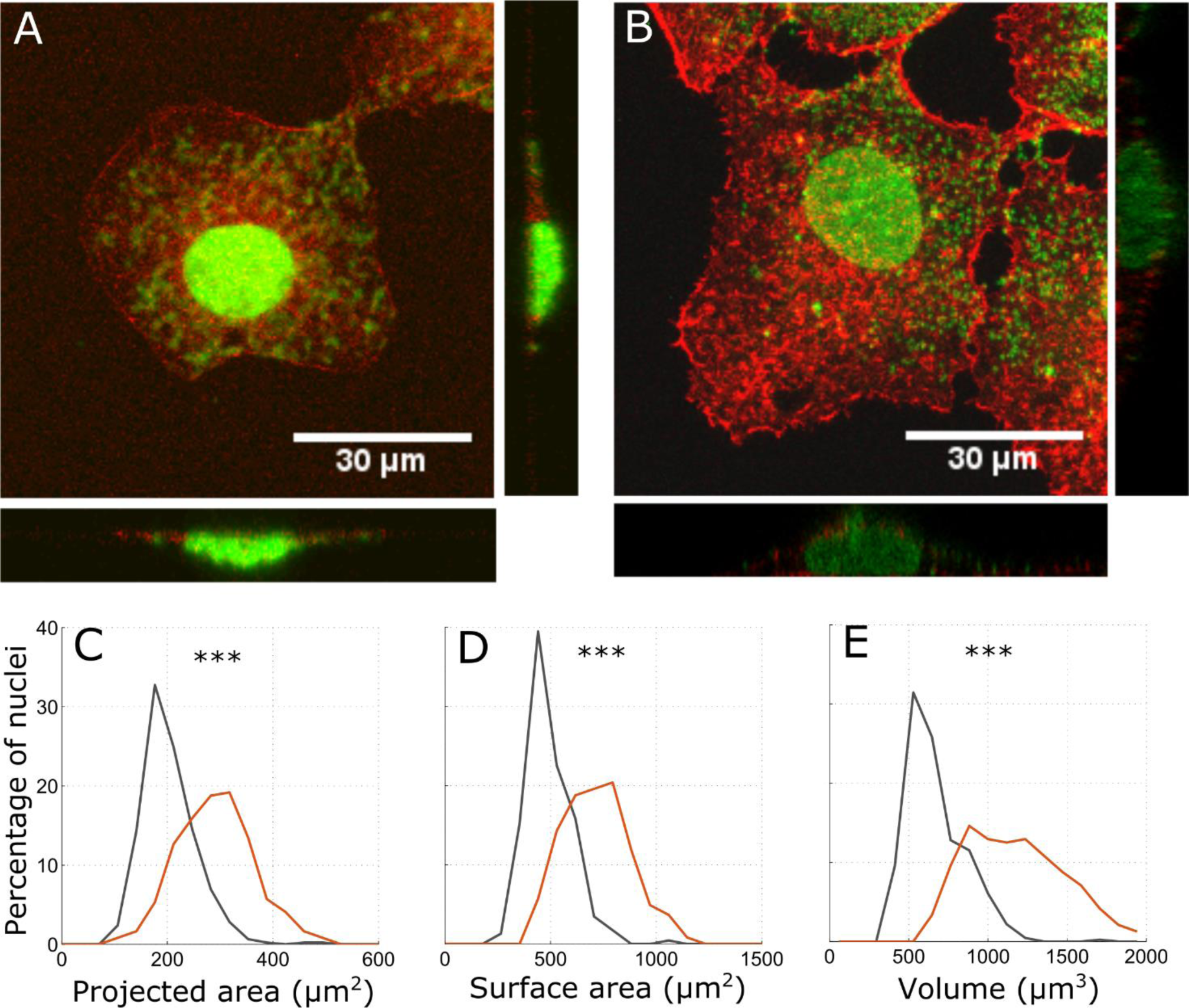
Morphology of the nucleus of Huh7 and HCV Replicon cells – Confocal images of Huh7 (A) and HCV Replicon cells (B). Actin is stained in red and nucleus in green. Probability distribution of projected area (C) surface area (D) and volume (E) of the nuclei of Huh7 (black) (N = 461) and HCV replicon (red) (N = 246) cells. Projected area, surface area and volume of the nuclei of HCV replicon cells are significantly larger than the nuclei of Huh7 cells. *** indicates p < 0.001 by two-tailed Kolmogrov-Smirnov test.

**Fig. 3.**
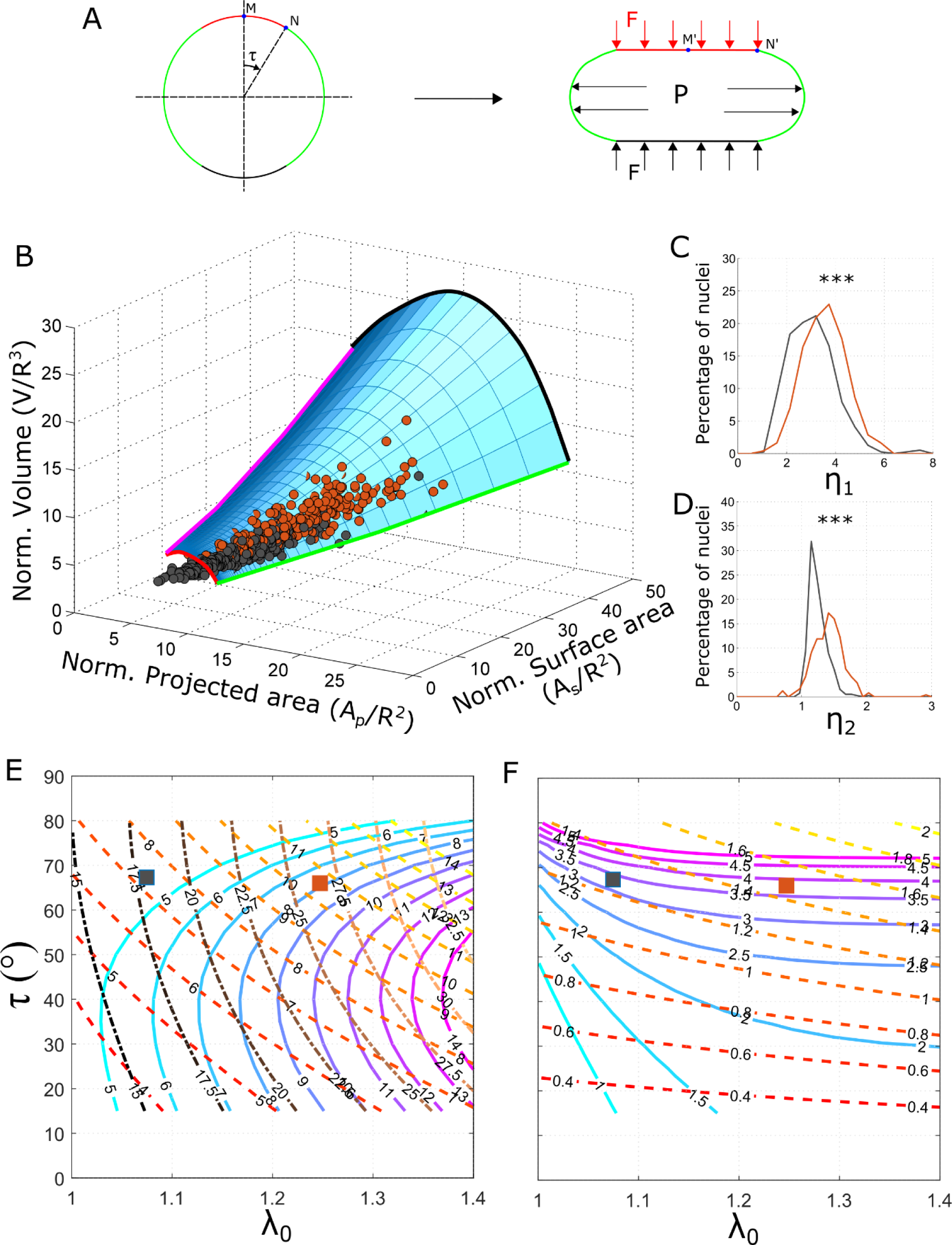
Non-dimensional mechanical model of nuclear morphology – (A) The spherical nuclear envelope deforms from a sphere by two forces: (i) an inflating pressure *P* and (ii) a force *F* due to cortical actin. The force from cortical actin is assumed to be arising from a flat plate that is pushing down on the nucleus. Contact between actin and nuclear envelope in the deformed state is along the red, horizontal line on the top of the nucleus. The corresponding region in the undeformed configuration is also marked in red. Points N and N’ are the boundary of the contact region in the deformed and undeformed configurations respectively. The angle subtended by N with the axis of symmetry is the contact angle τ. The stretch at the apex point of the nucleus, M’, is λ_0_. (B) Blue surface represents a relationship between the normalized projected area, surface area and volume of individual nuclei as predicted by the model. Black and red dots are the experimentally measured morphologies of the nuclei of Huh7 and HCV replicon cells. Almost all individual nuclei lie on the surface predicted by the model. The different boundaries of the surface are marked. Red and black curves are the lower and higher limits respectively of the initial stretch *λ*_*0*_. Magenta and green curves are the lower and higher limits, respectively, of the contact angle *τ*. Probability distributions of the non-dimensional parameters *η*_1_ (C) and *η*_2_(D) obtained from the experimentally measured nuclear morphologies (Fig. 2). *η*_1_ and *η*_2_ are significantly larger for HCV replicon cells (red) in comparison to Huh7 cells (black). *** p < 0.001 by two–tailed Kolmogorov – Smirnov test. (E) Contour curves of normalized volume (solid lines colored blue to magenta), normalized projected area (dashed lines colored red to yellow) and normalized surface area (dash-dot lines colored dark to light brown) as a function of *λ*_*0*_ and *τ* (F) Contour curves of *η*_1_ (solid lines colored blue tomagenta) and *η*_2_ (dashed lines colored red to yellow) as a function of *λ*_*0*_ and *τ*. The values of *η*_1_ and *η*_2_ for any nucleus can be obtained using these contour plots. From the volume, projected area and surface area of a nucleus, we calculate the normalized volume, projected area and surface area using Eq. (2.2) by assuming *R*. Using the normalized nuclear shape parameters in (E), obtain *λ*_*0*_ and *τ*. Using *λ*_*0*_ and *τ*, obtain *η*_1_ and *η*_2_ from (F). To illustrate the method, we have plotted the mean nuclear morphology of Huh7 (black square) and HCV replicon (red square) cells on (E) and (F).

For given *λ*_0_ and *τ*, we first calculated *η*_1_ and *η*_2_, and then numerically integrated the governing equations to obtain the principal strains. From the principal strains, we obtained normalized nuclear morphology, which is the deformed shape when a spherical membrane of unit radius is deformed by *η*_1_ and *η*_2_. The normalized nuclear morphology was characterized by normalized nuclear shape parameters, namely, projected area (*a*_*p*_), surface area (*a*_*s*_) and volume (*v*), defined earlier. To obtain the actual nuclear morphology, we scaled the normalized nuclear morphology by *R*. The normalized nuclear shape parameters are related to the actual nuclear shape parameters through scaling relations.

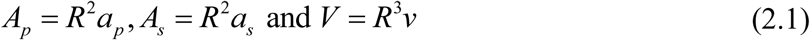

To obtain *η*_1_ and *η*_2_ corresponding to experimentally measured nuclei, we first normalized its nuclear shape parameters

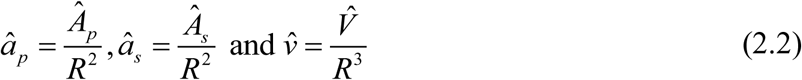

where ‘^’ denote experimentally measured values. We have assumed that our control and HCV-infected cells are descendant from a single clone and hence we have assumed the same *R* for all nuclei. *η*_1_ and *η*_2_ corresponding to each nucleus were then obtained using a least squares minimization by solving the following problem

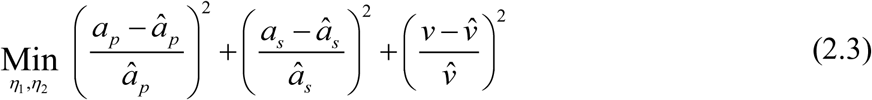

## Results

### HCV proteins alter the morphology of the nucleus

We measured the morphology of the nuclei of Huh7 and HCV replicon cells from confocal images. We obtained the boundary surface of the nucleus from confocal images using a 3D image processing algorithm (see supplementary information and Supplementary Movie 1). From the boundary surface, nuclear shape parameters such as projected area, surface area and volume were calculated. The nuclear shape parameters of Huh7 and HCV replicon cells are shown in Fig. 2 *C-E*. As seen in Figure 2, HCV replicon cells show increased projected area, surface area and the volume of the nucleus. The mean projected area of the nuclei in HCV replicon cells is equal to 312 μm^2^ whereas that of its control cell line (Huh7) is 220 μm^2^. The mean projected area of HCV replicon cells is approximately 42% higher than that of the control cells. Nuclei of HCV replicon cells have a mean surface area of 768 μm^2^ which is 44% higher than the mean surface area of control cells (534 μm^2^). The mean volume of the nucleus of HCV replicon cells (1.218×10^3^ μm^3^) is 88 % larger than that of control cells (0.667×10^3^ μm^3^).

By using the model, we obtained *η*_1_ and *η*_2_ corresponding to individual nuclei. The differences in *η*_1_ and *η*_2_ between Huh7 and HCV replicon cells were used to identify the reasons for the change in nuclear morphology.

### Two-parameter, non-dimensional mechanical model

In our computations we systematically varied *λ*_0_ and *τ* to obtain the corresponding normalized projected area (*a*_*p*_), surface area (*a*_*s*_) and volume (*v*) corresponding to each of them. These normalized nuclear shape parameters form a surface in *a*_*p*_ - *a*_*s*_*- v* space. This surface (blue surface in Fig. 3*B*) represents a relation among *a*_*p*_, *a*_*s*_ and *v* as predicted by the model. The bounds of the surface (blue surface in Fig. 3*B*) are due to limits on the simulation parameters. Since the nuclear envelope is assumed to be a membrane, it cannot resist compression. This sets a lower bound on *λ*_0_ (red boundary, *λ*_0_ *=* 1.001). Even though there is no upper bound for *λ*_0_, we have simulated only up to *λ*_0_ *=* 1.4, which was enough to contain all experimentally measured nuclear morphologies (black curve). The lower and higher limits of *τ* are 0° and 90° respectively. However, we have simulated from *τ =* 15° (magenta curve) to *τ =* 80° (green curve).

By assuming *R* = 5.5 µm, we obtained 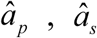 and 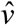 (Eq. (2.2)) for all experimentally measured nuclear morphologies (Fig. 2 *C-E*). 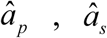 and 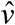 from individual, experimentally measured nuclei (black dots are Huh7 cells and red dots are HCV replicon cells in Fig. 3*B*), lie on the surface predicted by the model (Fig. 3*B* and Supplementary Movie 2). Since the nuclei in this study have low aspect ratio (Fig. 2 *A* and *B*, height to diameter around 0.18) they lie on the model surface in regions of large *τ* (between 55° and 75°). The nuclear morphologies would lie on the surface even if we change the radius in the unstressed state (*R* = 5 µm and 6 µm are shown in Fig. S5). At low values of *R*, more points will lie on the surface. However, many nuclei (larger 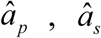 and 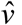) will have large strains (*λ* > 2). At high values of *R*, many nuclei were below the lower bound on *λ*_0_ (red boundary of the blue surface in Fig. 3*D* and Fig. S5*D*). Hence, we used an intermediate value of *R* = 5.5 µm. Nuclear morphology of HeLa cells obtained from (22) also lie on the surface predicted by our model (see Supplementary Movie 3).

We fit the model to 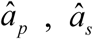 and 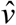 of Huh7 and HCV replicon cells, using Eq. (2.3), and obtained *η*_1_ and *η*_2_. Only those fits with relative errors < 0.05 (relative error in volume is defined as err_*v*_ *=* | (*v* − *v*’)/*v*’ | and similarly for projected area and surface area) for each of the normalized shape parameters are plotted in Fig. 3*C*. With an upper limit in error of 5%, we were able to fit around 85% of Huh7 nuclei and 99% of HCV replicon nuclei. We observed that both *η*_1_ and *η*_2_ are significantly larger for HCV replicon cells in comparison to Huh7 cells (p < 0.001 using two-tailed Kolmogrov-Smirnov test). The mean value of *η*_1_ is 3.33 for Huh7 cells and 3.81 for HCV replicon cells. The mean value of *η*_2_ is 1.29 for Huh7 cells and 1.45 for HCV replicon cells. Increase in *η*_1_ suggests that the nuclear envelope of HCV replicon cells have a lower elastic modulus in comparison to Huh7 cells. Increase in *η*_2_ suggests that the pre-tension in cortical actin is higher in HCV replicon cells in comparison to Huh7 cells. These predictions do not change for R = 5 µm and 6 µm (Fig. S5). Furthermore, *η*_1_ and *η*_2_ are similar to those *F Pπ R*2 estimated from individual values of the forces, material properties and initial geometry in (11): *PR*/μ*H =* 3.9 and 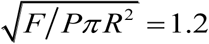

The values of *η*_1_ and *η*_2_ can be approximately estimated using a graphical method instead of solving the minimization problem in Eq. (2.3). *λ*_0_ and *τ* corresponding to a nuclear morphology can be obtained by locating 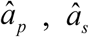 and 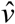 on the contour plot shown in Fig. 3*E*. *η*_1_ and *η*_2_ can then be obtained from the corresponding point in the contour plot shown in Fig. 3*F*. *η*_1_ and *η*_2_ for the mean nuclear morphology of Huh7 and HCV replicon cells were obtained using this technique (Fig. 3 *E* and *F*). Alternatively, these steps can be done numerically using the MATLAB codes provided with the Supplementary Data.

### HCV proteins increase cell stiffness and decrease nuclear stiffness

#### Cell stiffness

The stiffness of adherent cells, measured using AFM, is due to pre-tension in cortical actin, microtubules and the nucleus. However, the major contributing factor is the pretension in cortical actin (up to 50%) (15). Hence, we used cell stiffness measured using AFM as an indicator of the change in pre-tension in cortical actin. For comparing the cell stiffness between Huh7 and HCV replicon cells, we used the apparent modulus of elasticity obtained by fitting a suitable contact model to the force-displacement curves (23) (Supplementary information, Fig. S6). The elastic modulus of HCV replicon cells (263 ± 157 Pa) was found to be higher than that of Huh7 cells (203 ± 134 Pa) (Fig. 4*E*) but the increase was not statistically significant. The mean stiffness of HCV replicon cells was around 25% higher than that of Huh7 cells. The increase in stiffness suggests higher pre-tension in actin in HCV replicon cells in comparison to Huh7 cells.

**Fig. 4.**
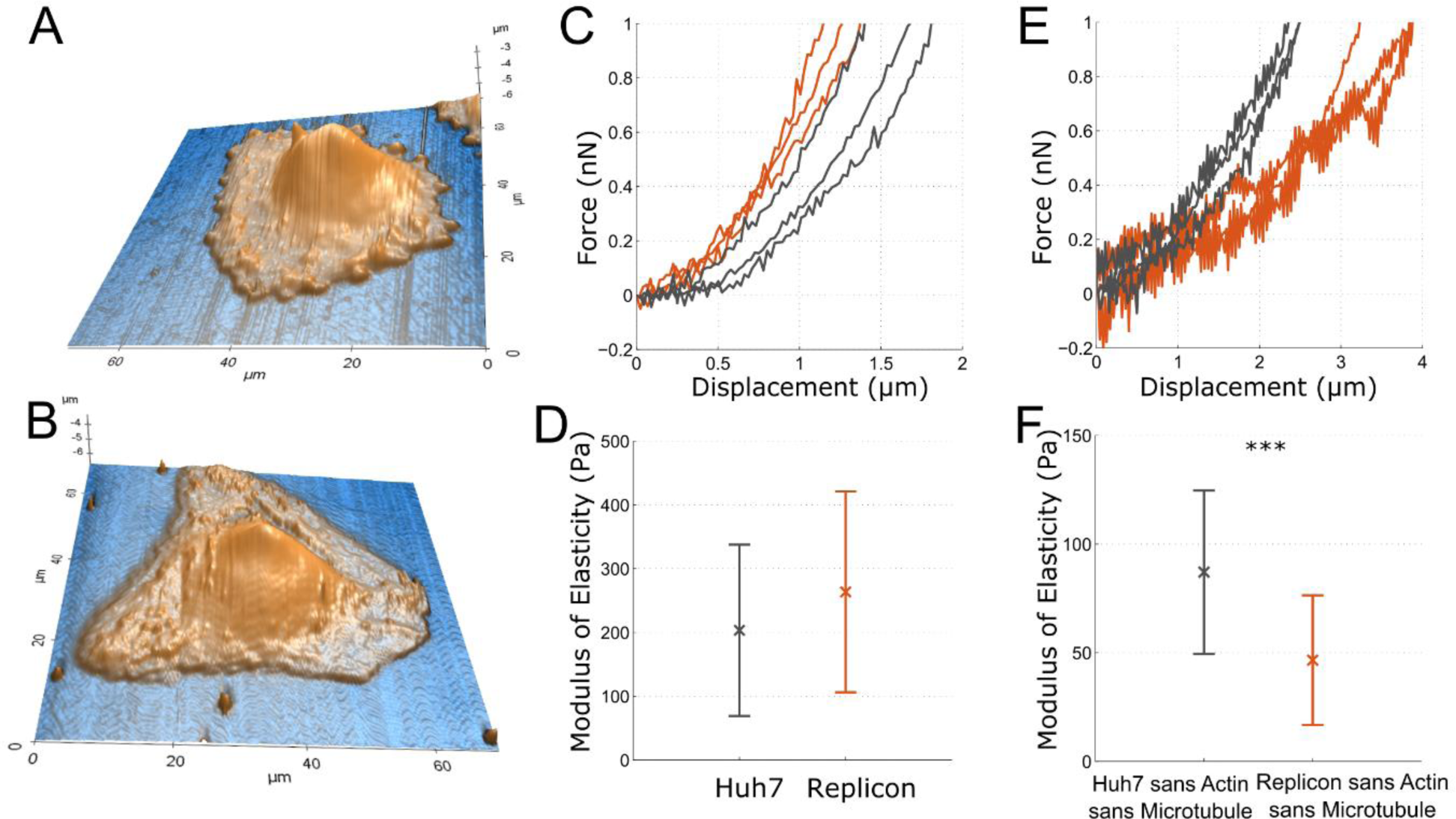
Mechanical characterization of Huh7 and HCV Replicon cells using AFM – Topography of Huh7 (A) and HCV Replicon cells (B) by contact mode imaging. Sample F-d curves (C) and apparent elastic modulus (D) of Huh7 (black, N = 21) and HCV Replicon (red, N = 23) cells. Sample F-d curves (E) and apparent modulus of elasticity (F) of Huh7 cells (black, N = 34) and HCV replicon cells (red, N = 20) with actin and microtubule depolymerised. *** p < 0.001 by one–tailed Student’s t-test

**Fig. 4.**
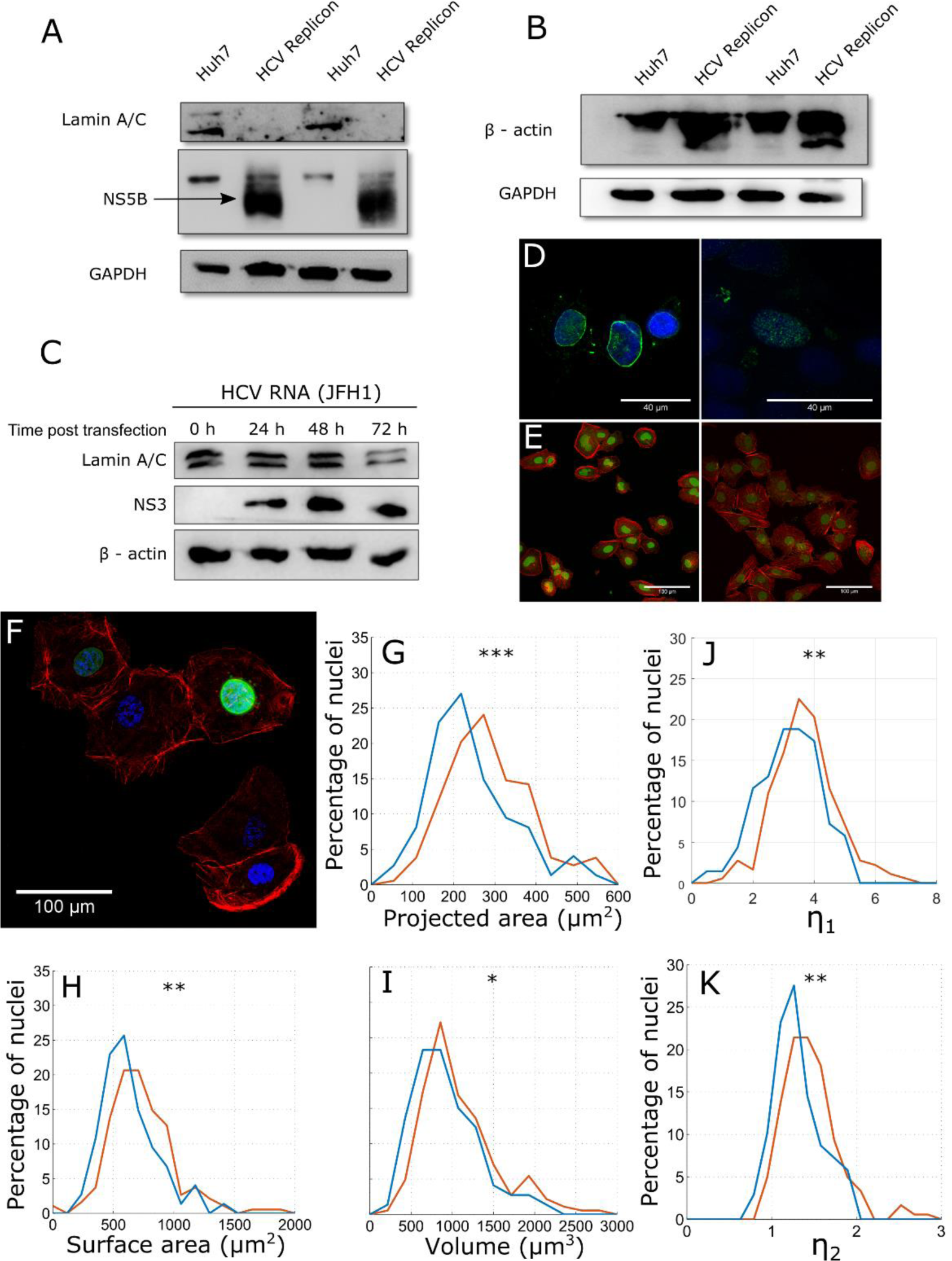
Downregulation of lamin-A,C by HCV – (A) lamin-A,C levels in HCV replicon cells, (B) β-actin levels in HCV replicon cells (C) Change in lamin-A,C level with time under HCV RNA transfection. (D) Immuno-fluorescence of Huh7 cells (left) and Huh7 cells transfected with HCV RNA. Nucleus is stained in blue by DAPI and lamin-A,C in green (E) Immuno-fluorescence of Huh7 cells (left) and HCV replicon cells (right). Nucleus is stained in green and actin in red. (F) HCV replicon cells tranfected with lamin-A–GFP over-expression plasmid. Nucleus is stained in blue, actin in red and lamin-A–GFP in green. Probability distribution of the projected area (G), surface area (H) and volume (I) of the nuclei of HCV replicon (red) (N = 192) and HCV replicon cells expressing lamin-A,C-GFP (blue) (N = 74). Probability distributions of the non-dimensional parameters *η*_1_(J) and *η*_2_(K) obtained from the nuclear morphologies in G-I. *η*_1_ and *η*_2_ are significantly larger for HCVreplicon cells (red) in comparison to those over-expressing lamin-A (blue). * p < 0.05, ** p < 0.01 and *** p < 0.001 by two–tailed Kolmogrov–Smirnov test.

#### Nuclear stiffness

A recent study has shown that the nucleus becomes the major load-bearing member when the actin and microtubules are de-polymerized. The authors observed that while indenting using an AFM tip, the nuclear deformation increased from 5% to 30% upon depolymerization of the actin cytoskeleton and microtubules (24). This suggests that we can obtain the stiffness of the nucleus by measuring the cell stiffness after depolymerizing the actin cytoskeleton and microtubules. Hence, we used Cytochalasin D and Nocadazole to depolymerize actin and microtubule respectively, and measured the stiffness using AFM. The stiffness of the nuclei of HCV replicon cells was significantly lower (p < 0.001) than that of Huh7 cells (Fig. 4*F*). Nuclei of HCV replicon cells had a mean apparent elastic modulus of 49 ± 26 Pa while those of Huh7 cells was 87 ± 37 Pa. The reduction in nuclear stiffness is around 43%.

Our results suggest that HCV proteins marginally increase the stiffness of the cells but significantly reduce the stiffness of the nuclei. The stiffness of the cells was measured right above the nucleus (see Supplementary information and Fig. S6) and hence the measured stiffness will have contributions from the cortical actin as well as the nucleus. Hence, the increase of prestress in actin could be masked by a reduction in stiffness of the nucleus. This could be the reason why we do not see a significant increase in the stiffness of HCV replicon cells in comparison to Huh7 cells.

### Hepatitis C Viral proteins down-regulate lamin-A,C and up-regulate actin

#### Down-regulation of lamin-A,C

We measured the expression levels of lamin-A,C and actin for biochemical confirmation of the predictions from the computational model. We observed that Hepatitis C Virus down-regulates lamin-A,C in Huh7 cells. The down-regulation of lamin-A,C was observed in HCV replicon cells as well as those transfected with full-length HCV RNA. The down-regulation of lamin-A,C was confirmed by western blot (Fig. 5 *A* and *C*) as well as immunofluorescence (Fig. 5*D*). In Huh7 cells, lamin-A,C is localized at the nuclear periphery whereas in Huh7 cells transfected with HCV RNA, the levels of lamin-A,C were down-regulated and do not localize to the nuclear periphery (Fig. 5*D*). However, there was no difference in the expression level of lamin-B1 between Huh7 and HCV Replicon cells (Fig. S1*C*). We further checked the levels of lamin-A,C in Coxsackie Virus B3, which is another single stranded positive sense RNA Virus. However, upon Coxsackie viral infection, lamin-A,C levels were not down-regulated (Fig. S1*D*). Hence, lamin-A,C down-regulation may be a specific strategy of HCV and not a general mechanism of cytoplasmic RNA viruses.

To establish the causal relationship between the changes in nuclear mechanics and down-regulation of lamin-A,C, we over-expressed lamin-A in HCV replicon cells and measured the effect on nuclear morphology. We used a lamin-A-GFP construct so that the cells that were over-expressing lamin-A could be identified by fluorescence imaging (Fig. 5*F*). We observed that the nuclei of HCV replicon cells over-expressing lamin-A had lower projected area, surface area and volume as compared to the nuclei of HCV replicon cells (Fig. 5 *G-I*). The mean projected area of the nuclei of HCV replicon cells over-expressing lamin-A was 271 µm^2^ while that of the nuclei of HCV replicon cells was 330 µm^2^. The mean surface area of the nuclei of HCV replicon cells over-expressing lamin-A was 686 µm^2^ while that of the nuclei of HCV replicon cells was 805 µm^2^. The mean volume of the nuclei of HCV replicon cells over-expressing lamin-A was 1048 µm^3^ while that of the nuclei of HCV replicon cells was 1255 µm^3^. Rescue of nuclear morphology upon over-expressing lamin-A confirms that the changes in nuclear mechanics in HCV replicon cells are due to downregulation of lamin-A,C.

#### Up-regulation of actin

HCV replicon cells showed higher expression of actin in comparison to Huh7 cells. This was confirmed both by western blot (Fig. 5*B*) as well as immuno-fluorescence (Fig. 5*E*). However, actin up-regulation was not observed under HCV RNA transfection (Fig. 5*C*). This could be because the up-regulation of actin is a long-term effect of HCV non-structural proteins and is hence not observed in a short-term, transient transfection of HCV RNA. In contrast, we did not observe any significant difference in the expression of microtubules between Huh7 and HCV replicon cells (Fig. S1 *A* and *B*).

#### Interplay between lamin-A,C and *E*_*1*_

Since over-expressing lamin-A increases the modulus of the nuclear envelope, the model predicts that *η*_1_ will be lower for HCV replicon cells over-expressing lamin-A in comparison to HCV replicon cells. To validate this, we calculated *η*_1_ and *η*_2_ from the nuclear morphology of HCV replicon cells and those over-expressing lamin-A. We observed that the HCV replicon cells over-expressing lamin-A (mean value of *η*_1_= 3.45) have significantly lower *η*_1_ in comparison to HCV replicon cells (mean value of *η*_1_= 3.97). The reduction in the size of the nucleus and *η*_1_ upon over-expressing lamin-A further validates our model. Interestingly, *η*_2_ is also lower in HCV replicon cells over-expressing lamin-A in comparison to HCV replicon cells.

Even though our experimental results, using AFM, western blot and immunofluorescence, confirm decrease in the modulus of the nuclear envelope and an increase in the pre-tension in cortical actin, the possibility of change in the inflating pressure cannot be discounted. Depolymerization of microtubules can increase the net inflating pressure (11) and hence increase *η*_1_. However, we have not observed any difference in the expression of microtubules between Huh7 and HCV replicon cells (Fig. S1 *A* and *B*). Another factor that can increase the inflating pressure is an increase in DNA content or de-condensation of chromatin. Previous studies have established cell cycle arrest at S (25) and G2 (26) phases in HCV infection, which could increase the DNA content inside the nucleus. Hence alterations in the DNA content could be another contributing factor for the changes in the morphology of the nucleus.

## Discussion

We presented a non-dimensional, two-parameter elastic model that gives morphological parameters (volume, surface area and projected area) of the nucleus in its native adherent state in terms of net inflation pressure and force due to cortical actin. There exists a relationship among the three morphological parameters, which is depicted as a surface in Fig. 3*B*. Experimentally measured volume, surface area and projected area of individual nuclei lie on this surface. This applies to our data and those reported in (22). Every point on the surface is associated with two non-dimensional parameters expressed in terms of the inflation pressure and force due to cortical actin as well as elastic modulus, radius and thickness of the nuclear envelope. For known values of the radius and thickness of the nuclear envelope, the first non-dimensional parameter gives the ratio of inflation pressure to the elastic modulus and the second gives the ratio of the force due to cortical actin and the inflation pressure as per equations shown in Fig. 1. (See also Eqs. (3.29) and (3.35) in the Supplementary Information). The two non-dimensional parameters together also give the ratio of cortical actin force to modulus of the nuclear envelope.

It may be noted that the non-dimensional nuclear parameters can be approximately estimated using a graphical method or a numerical method (See Supplementary Data for the numerical method). Hence, our model can be used to study the effect of stimuli on the nuclear envelope and cytoskeleton by just measuring the nuclear morphology. For example, a recent study (27) showed that increased substrate stiffness concomitantly increased the expression of lamin-A,C and myosin in mesenchymal stem cells thereby increasing the modulus of the nuclear envelope and the pretension in actin cytoskeleton. Using our model, these results could be inferred from the nuclear morphology by a decrease in *η*_1_Fig. 1). and an increase in *η*_2_ (see equations in Fig. 1).

Our model is axisymmetric and hence does not consider the eccentricity of the nucleus. For the cells used in this study, the eccentricity of the nucleus is around 0.6 – 0.7, which translates to a low aspect ratio (major axis to minor axis) of 1.25 – 1.4. Our model may not be applicable to cells with elongated nuclei (aspect ratio > 2) such as fibroblasts. However, since the model predicts a relationship among the projected area, surface area and volume of individual nuclei (blue surface in Fig. 3*B*), it can be used to verify the applicability of our model to nuclei of any cell type. MATLAB codes and modeling data for plotting individual nuclear morphology over the model surface are provided (Supplementary Data) for verification.

The inflating pressure in our equations is the net effect of the osmotic pressure, and forces from chromatin and microtubules. Therefore, an indication of change in pressure (increase in *η*_1_ and decrease in *η*_2_ or vice versa) cannot be attributed to any of these without further experimental measurements.

The force exerted by cortical actin on the nucleus is a function of the pre-tension in cortical actin and the contact area between the nuclear envelope and cortical actin. The contact area in turn depends on the inflating pressure. Thus, the contact force is not independent of the inflating pressure. Hence, there could be a relation between *η*_1_ and *η*_2_. We have observed a positive correlation between *η*_1_ and *η*_2_ for Huh7 and HCV replicon cells. Pearson’s correlation coefficient for Huh7 cells was 0.96 and that for HCV replicon cells was 0.88. However, a part of this correlation could be due to dependencies between the parameters in *η*_1_ and *η*_2_ arising from biochemical mechanisms. For example, expression of lamin-A,C (*E*_1_) is known to increase with pre-tension in cortical actin (*F*) in mesenchymal stem cells (27).

We showed the applicability of our two-parameter model in the case of HCV replicon cells in which lamin-A,C is down-regulated and cortical actin is up-regulated. It is interesting that HCV, which is a cytoplasmic RNA virus, affects nuclear mechanics. Such phenomena are well known in the case of DNA viruses such as Herpes Simplex Virus (28), HIV (29) and SV40 (30) wherein they disrupt the nuclear lamina by down-regulating lamin-A,C or B. The replication complexes of these viruses are assembled inside the nucleus. They need to enter the nucleus and export the viral particles produced to the cytoplasm. The nuclear lamina forms a barrier to these transports and hence these viruses need to disassemble the nuclear lamina. But such considerations are absent in the case of RNA viruses such as HCV since their entire life cycle is confined to the cytoplasm. However, our study shows that HCV is disrupting the nuclear lamina by down-regulating lamin-A,C and thereby deregulating important functions of the nucleus. Apart from its role in gene-regulation, lamins are shown to be mechano-sensors (31). They are known to translate mechanical stimuli from the exterior of the cell into appropriate changes in gene expression by forming a mechanical signaling pathway through the cytoskeleton and LINC (Linker of Nucleoskeleton to Cytoskeleton) to the interior of the nucleus (32). Hence, by disrupting the nuclear lamina, HCV could be impairing the mechanobiological homeostasis of liver cells.

In summary, we have related morphological, biochemical and biomechanical measurements to the physical properties of the nucleus and the cell through a mechanical model. The model predicts a relationship among the projected area, surface area and volume of individual nuclei. These morphological parameters can be easily obtained by confocal imaging and the relationship can then be used to ascertain the applicability of our model to any cell type. Once the model is found to be suitable, the non-dimensional parameters corresponding to individual nuclei can be estimated. The changes in the non-dimensional parameters due to a stimulus suggest perturbations in the nuclear envelope and cytoskeleton, which can then be used to guide further experimental studies. We have used this technique in liver cells and discovered alterations in nuclear mechanics due to Hepatitis C Virus.

## Acknowledgments

The authors would like to thank Monisha Mohandas for help with AFM experiments; Anoosha Pai and Supriya M.V for help with Finite Element Analysis; Dr. Biju George for Coxsackie Virus infection; Department of Biotechnology, Govt. of India for funding; and Prof. G. V. Shivashankar and Prof. Dennis Discher for useful comments.

## References

1. Pajerowski JD, Dahl KN, Zhong FL, Sammak PJ, Discher DE (2007) Physical plasticity of the nucleus in stem cell differentiation. Proc Natl Acad Sci U S A 104(40).

2. Cao X, et al. (2016) A Chemomechanical Model for Nuclear Morphology and Stresses during Cell Transendothelial Migration. Biophys J 111(7):1541–1552.

3. Denais CM, et al. (2016) Nuclear envelope rupture and repair during cancer cell migration. Science (80-) 352(6283):353–358.

4. Makhija E, Jokhun DS, Shivashankar G V (2015) Nuclear deformability and telomere dynamics are regulated by cell geometric constraints. Proc Natl Acad Sci U S A 113(1):E32–40.

5. Irianto J, Pfeifer CR, Ivanovska IL, Swift J, Discher DE (2016) Nuclear Lamins in Cancer. Cell Mol Bioeng 9(2):258–267.

6. De las Heras JI, Batrakou DG, Schirmer EC (2013) Cancer biology and the nuclear envelope: A convoluted relationship. Semin Cancer Biol 23(2):125–137.

7. Lammerding J, et al. (2004) Lamin A / C deficiency causes defective nuclear mechanics and mechanotransduction. J Clin Invest 113(3):370–378.

8. Worman HJ, Ostlund C, Wang Y (2010) Diseases of the nuclear envelope. Cold Spring Harb Perspect Biol 2(2):1–17.

9. Folker ES, Ostlund C, Luxton GWG, Worman HJ, Gundersen GG (2011) Lamin A variants that cause striated muscle disease are defective in anchoring transmembrane actin-associated nuclear lines for nuclear movement. Proc Natl Acad Sci 108(1):131–136.

10. Zwerger M, et al. (2013) Myopathic lamin mutations impair nuclear stability in cells and tissue and disrupt nucleo-cytoskeletal coupling. Hum Mol Genet 22(12):2335–2349.

11. Kim D-H, et al. (2015) Volume regulation and shape bifurcation in the cell nucleus. J Cell Sci 128(18):3375–85.

12. Feng WW, Yang WH (1973) On the Contact Problem of an Inflated Spherical Nonlinear Membrane. J Appl Mech 45(1):209–214.

13. Schape J, Praube S, Radmacher M, Stick R (2009) Influence of lamin A on the mechanical properties of amphibian oocyte nuclei measured by atomic force microscopy. Biophys J 96(10):4319–4325.

14. Lammerding J, et al. (2006) Lamins A and C but Not Lamin B1 Regulate Nuclear Mechanics. J Biol Chem 281(35):25768–25780.

15. Barreto S, Clausen CH, Perrault CM, Fletcher DA, Lacroix D (2013) A multi-structural single cell model of force-induced interactions of cytoskeletal components. Biomaterials 34(26):6119–6126.

16. Lohmann V, et al. (1999) Replication of Subgenomic Hepatitis C Virus RNAs in a Hepatoma Cell Line. Science (80-) 285(5424):110–113.

17. Finan JD, Chalut KJ, Wax A, Guilak F (2009) Nonlinear Osmotic Properties of the Cell Nucleus. Ann Biomed Eng 37(3):477–491.

18. Finan JD, Guilak F (2009) The effects of osmotic stress on the structure and function of the cell nucleus. J Cell Biochem 31(9):n/a–n/a.

19. Cao X, et al. (2016) A Chemomechanical Model for Nuclear Morphology and Stresses during Cell Transendothelial Migration. Biophys J 111(7):1541–1552.

20. Rowat AC, Lammerding J, Ipsen JH (2006) Mechanical properties of the cell nucleus and the effect of emerin deficiency. Biophys J 91(12):4649–4664.

21. Green AE, Adkins JW (1960) Large elastic deformations (The Clarendon Press, Oxford).

22. Velliste M, Murphy RF (2002) Automated determination of protein subcellular locations from 3D fluorescence microscope images. Biomedical Imaging, 2002. Proceedings. 2002 IEEE International Symposium On, pp 867–870.

23. Johnson KL (1987) Contact Mechanics (Cambridge university press).

24. Haase K, et al. (2016) Extracellular Forces Cause the Nucleus to Deform in a Highly Controlled Anisotropic Manner. Sci Rep (October 2015):1–11.

25. Yang XJ, et al. (2006) HCV NS2 protein inhibits cell proliferation and induces cell cycle arrest in the S-phase in mammalian cells through down-regulation of cyclin A expression. Virus Res 121(2):134–143.

26. Kannan RP, Hensley LL, Evers LE, Lemon SM, McGivern DR (2011) Hepatitis C Virus Infection Causes Cell Cycle Arrest at the Level of Initiation of Mitosis. J Virol 85(16):7989–8001.

27. Buxboim A, et al. (2014) Matrix elasticity regulates lamin-A,C phosphorylation and turnover with feedback to actomyosin. Curr Biol 24(16):1909–17.

28. Scott E, O’Hare P (2001) Fate of the inner nuclear membrane protein lamin B receptor and nuclear lamins in herpes simplex virus type 1 infection. J Virol 75(18):8818–8830.

29. de Noronha CM, et al. (2001) Dynamic disruptions in nuclear envelope architecture and integrity induced by HIV-1 Vpr. Science 294(5544):1105–1108.

30. Butin-Israeli V, et al. (2011) Simian virus 40 induces lamin A/C fluctuations and nuclear envelope deformation during cell entry. Nucleus 2(4):320–330.

31. Swift J, et al. (2013) Nuclear Lamin-A Scales with Tissue Stiffness and Enhances Matrix-Directed Differentiation. Science (80-) 341(6149):1240104–1240104.

32. Isermann P, Lammerding J (2013) Nuclear mechanics and mechanotransduction in health and disease. Curr Biol 23(24):1113–1121.

